# Less is more: Hormetic and selective antimicrobial activity of a thymoquinone-standardized black seed oil ThymoQuin^®^ in gut microbiota models

**DOI:** 10.64898/2026.05.10.724082

**Authors:** Flavia Stal Papini, Nadine Zettner, Samar Sawas, Cornelius Roth, Lisa Bäumer

## Abstract

The gut microbiome plays a central role in host metabolism, immune function, and overall health, with disruptions in microbial composition (dysbiosis) being associated with a range of metabolic, inflammatory, and infectious conditions [1,2]. Consequently, strategies aiming to modulate the microbiome require selective activity that preserves beneficial commensals while limiting pathogenic organisms [3]. In this context, ThymoQuin^®^—a cold-pressed, standardized black cumin (*Nigella sativa*) seed oil developed by TriNutra Ltd. and defined by ≥3% thymoquinone (TQ), controlled p-cymene levels, and low free fatty acids (≤1.25%)—was evaluated for its microbiome-relevant activity. *In vitro* minimum bactericidal concentration (MBC) assays across three independent batches demonstrated a biphasic, dose-dependent response. At intermediate concentrations (0.25–0.5%), *Streptococcus thermophilus* was strongly stimulated (up to 53-fold) and *Lactiplantibacillus plantarum* fully preserved, while *Klebsiella pneumoniae* was effectively reduced (>94%). *Akkermansia muciniphila* exhibited stable viability at concentrations below 1%, with reductions only observed at 1%. This is notable given its role as a mucin-degrading commensal that has been linked to metabolic health, but whose abundance may vary across physiological and disease contexts [4,5]. At concentrations ≥1%, selective effects diminished, resulting in broader antimicrobial activity and reduced specificity. These findings indicate a defined concentration range in which selective microbiome modulation is maintained, whereas higher thymoquinone levels may increase the risk of non-selective detrimental effect on microbes.

## 1. Introduction

The human gut microbiome is a complex and functionally diverse microbial ecosystem that contributes to host physiology through multiple complementary mechanisms, including digestion of otherwise indigestible substrates, production of short-chain fatty acids (SCFAs), maintenance of epithelial barrier integrity, colonization resistance against pathogens, modulation of mucosal and systemic immunity, and regulation of host metabolic homeostasis. These functions are not uniformly distributed across all commensal bacteria. Rather, different microbial taxa occupy distinct ecological and metabolic niches, and their contribution to intestinal homeostasis depends on their substrate preferences, oxygen tolerance, metabolic output, interaction with the mucus layer, and capacity to communicate with epithelial and immune cells. SCFAs, particularly acetate, propionate, and butyrate, are among the best-characterized microbiota-derived metabolites and have been implicated in epithelial energy metabolism, barrier regulation, and immune signaling [6]. Within this ecosystem, lactic acid bacteria, strict anaerobic butyrate producers, and mucin-associated organisms represent functionally distinct groups of particular relevance for gut health. Lactic acid bacteria are commonly associated with fermented foods and probiotic applications and may contribute to intestinal resilience through fermentation, production of antimicrobial metabolites, competitive exclusion of pathogens, and modulation of host responses [7, 8]. In contrast, obligate anaerobic butyrate-producing commensals are closely linked to colonic epithelial energy supply and anti-inflammatory mucosal homeostasis [9, 10]. Mucin-associated bacteria occupy a separate ecological niche at the host–microbe interface, where they participate in mucus-layer turnover and influence epithelial and immunometabolic signaling [11]. Therefore, when evaluating bioactive compounds with potential antimicrobial or microbiome-modulating activity, it is important to test their effects across taxonomically and functionally distinct gut-associated bacteria. Such an approach helps distinguish selective, microbiome-compatible activity from broad, non-specific antimicrobial inhibition [12]. There is growing scientific and commercial interest in natural compounds that selectively modulate gut microbiota: favoring beneficial bacteria, suppressing pathogens, and preserving the diversity of the microbial community.

Black cumin (*Nigella sativa*) seed represents a scientifically relevant candidate for microbiome-compatibility testing and has been used in traditional medicine for over 2,000 years across the Middle East, Southeast Asia, and North Africa [13]. Modern research has isolated thymoquinone (TQ) as the principal bioactive constituent, responsible for the oil’s well-documented antioxidant, anti-inflammatory, and antimicrobial properties [13, 14]. However, not all black seed oil products are equivalent — the market is characterized by enormous variation in TQ content, p-Cymene ratios, oil purity, and manufacturing quality.

ThymoQuin^®^, developed by TriNutra Ltd., represents a scientifically calibrated standard in black seed oil. Our study demonstrates how ThymoQuin^®^’s quality attributes and dose-response characteristics translate into optimal, selective gut microbiome performance — and why products with higher TQ concentrations may produce the opposite of the desired effect. In this context, a selection of key microbes present in the human gut were tested for their susceptibility towards ThymoQuin^®^ in the frame of this study. *Streptococcus thermophilus, Lactiplantibacillus plantarum, Klebsiella pneumoniae*, and *Akkermansia muciniphila* provides a biologically meaningful test panel to assess whether ThymoQuin^®^ exerts selective rather than broadly suppressive effects on representative beneficial or commensal gut-associated bacteria. These organisms cover complementary functional niches, including food-associated lactic acid fermentation, probiotic *Lactobacillaceae* activity, anaerobic butyrate production, and mucin-associated host–microbiota crosstalk.

*S. thermophilus* is a Gram-positive lactic acid bacterium widely used as a starter culture in yogurt and other fermented dairy products. Beyond its technological importance, it has gained interest as a beneficial bacterium because of reported health-relevant functions, including support of lactose digestion, production of exopolysaccharides, and other strain-dependent bioactive properties. Its long-standing use in food applications and its relevance to fermented-food-associated microbiota make it an appropriate organism for assessing whether preserves or supports beneficial lactic acid bacteria rather than inhibiting them non-selectively [15].

*L. plantarum* is a metabolically versatile lactic acid bacterium frequently investigated for probiotic potential. It is notable for its ability to survive diverse environmental conditions and for strain-dependent properties that may include antimicrobial activity, gastrointestinal persistence, modulation of immune responses, and support of epithelial barrier function. Including *L. plantarum* in the test panel is therefore relevant for determining whether ThymoQuin^®^ is compatible with a representative probiotic *Lactobacillaceae* species associated with gut resilience and pathogen exclusion [16].

*K. pneumoniae* was included as a clinically relevant opportunistic pathobiont and Gram-negative facultative anaerobe that can colonize the gastrointestinal tract. The gut is an important reservoir for *K. pneumoniae* and gastrointestinal carriage has been shown to contribute to subsequent infection risk, particularly in vulnerable or hospitalized patients. Therefore, inclusion of *K. pneumoniae* enabled assessment of whether ThymoQuin^®^ demonstrates selective inhibitory activity against an undesirable gut-associated pathobiont while preserving beneficial commensals involved in lactic acid fermentation, butyrate production, mucosal homeostasis, and host–microbiota signaling [17].

*A. muciniphila* is a mucin-degrading, mucosa-associated commensal that inhabits the intestinal mucus layer and participates in mucus turnover, epithelial barrier regulation, and host– microbiota communication. It has attracted considerable attention as a next-generation probiotic candidate, particularly in metabolic and inflammatory contexts. However, the biological role of *A. muciniphila* appears to be context-dependent: while many studies associate it with beneficial metabolic and barrier-related effects, excessive enrichment or disease-specific microbial configurations may not always be favorable. For this reason, testing *A. muciniphila* is particularly relevant, since unnecessary suppression of this taxon may be undesirable, whereas excessive or inappropriate modulation may also be biologically unfavorable depending on host and disease context [18].

Together, these four taxa provide a functionally differentiated model panel for assessing microbiome compatibility. By including lactic acid bacteria, a strict anaerobic butyrate producer and a mucin-associated commensal, the experimental design allows evaluation of whether ThymoQuin^®^ selectively affects specific microorganisms while preserving bacteria with recognized roles in gut homeostasis. This is particularly important for compounds derived from botanicals such as *N. sativa*, which may possess antimicrobial activity but should ideally avoid broad disruption of beneficial commensal taxa when positioned for gut-health applications.

## 2. Material and methods

### 2.1 Information on the test substance

ThymoQuin^®^, standardized black cumin seed oil (*N. sativa*, purity 3%) developed by TriNutra Ltd., was tested in three independent batches: 4816078, 4820084, and 4820086.

Each batch of the test substance was dissolved in squalane (99%, ThermoScientific, Lot #A0459480) to prepare a 10% (v/v) stock solution.

This concentration corresponds to a 10-fold stock relative to the highest final test concentration (1%). Subsequent 10-fold working stocks for lower concentrations (5%, 2.5%, etc.) were prepared by serial 2-fold dilutions of the 10% stock in squalane. The final squalane concentration in the tests was always 10%, independent of the product concentration tested.

### 2.2 Bacterial strains and growth conditions

**Table 1:** Bacterial species used in this study.

| Strains | Collection<br>Number | Liquid<br>Medium | Agar | Description | Reference /<br>Source |
| --- | --- | --- | --- | --- | --- |
| *Akkermansia*<br>*muciniphila* | DSM 22959 | Enriched<br>anaerobic<br>broth<br>(own<br>production) | Enriched<br>anarobic<br>agar<br>(own<br>production) | Anaerobic<br>strain,<br>isolated<br>from human<br>intestinal<br>tract | [19] |
| *Klebsiella*<br>*pneumoniae* | DSM<br>681 | CASO<br>(Carl Roth,<br>Karlsruhe,<br>Germany) | CASO | Aerobic<br>strain,<br>unknown<br>source | [20] |
| *Lactiplantibacillus*<br>*plantarum subs.*<br>*plantarum* | DSM 20174 | MRS<br>(Carl Roth,<br>Karlsruhe,<br>Germany) | MRS | Aerobic<br>strain,<br>isolated<br>from pickled<br>cabbage | [21, 22] |
| *Streptococcus*<br>*thermophilus* | DSM 20617 | M17<br>(Merck,<br>Darmstadt,<br>Germany) | M17 | Micro-<br>aerophilic<br>strain,<br>isolated<br>from<br>pasteurized<br>milk | [20] |

### 2.3 Minimal bactericidal concentration (MBC) assay

#### 2.3.1 Cultivation of the bacteria

Bacterial strains were pre-cultured from agar plates in appropriate growth media and incubated overnight at 37°C under species-specific conditions (including anaerobic conditions where required). Cells were harvested by centrifugation (4500 × g, 4°C, 5 min), washed in 1x phosphate-buffered saline (ROTI^®^Fair PBS 7.4, Carl Roth), and adjusted based on optical density (OD_600_=1). Standardized inoculi were prepared by diluting cultures in fresh, species-specific media to a defined starting density (OD_600_ = 0.1) for subsequent minimum bactericidal concentration (MBC) assays.

#### 2.3.2 MBC conditions

For the assays, sterile 12-well plates were prepared with a final volume of 2 mL per well. Treatment wells contained microbial suspension (1.8 mL) combined with either test substance dissolved in squalane or squalane alone as untreated control. ThymoQuin^®^ was tested at final concentrations of 0.031%, 0.063%, 0.125%, 0.25%, 0.5%, and 1%. Blank wells (medium with or without squalane, without inoculum) were included for each condition. Agar plates were incubated for 24 h at 37°C under anaerobic conditions (0% O_2_, 8% CO_2_, 2% H_2_).

Following incubation, samples were mixed and serially diluted in sterile PBS. Aliquots (100 µL) of appropriate dilutions were plated in triplicate onto agar plates and incubated under species-specific conditions. Colonies were counted on plates containing 30–300 colony-forming units (CFUs), and CFU/mL values were calculated based on dilution factors and plated volume.

#### 2.3.3 Statistical analysis

For each experimental condition, CFU counts were obtained from three replicate plates at the most appropriate dilution (i.e., the dilution yielding 30–300 colonies). If no dilution fell within this range, the nearest dilution with reliably countable colonies was selected.

Colony counts from each plate were converted to CFU/mL by multiplying the raw count by the corresponding dilution factor. For each organism and concentration, the mean CFU/mL and standard deviation (SD) were calculated from three technical replicates. The untreated control (squalane) mean and SD were calculated from at least six replicates and applied uniformly across all tested concentrations.

The log_10_ change relative to the untreated control was calculated as:

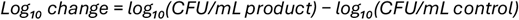

Negative values indicate inhibition of growth; positive values indicate an increase relative to the untreated control. The uncertainty of the log_10_ change was estimated by error propagation of the individual standard deviations:

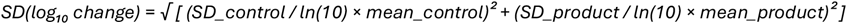

Results are additionally expressed as percentage of the untreated control, where the control is set to 100%:

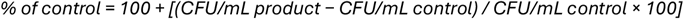

Values above 100% indicate growth relative to the untreated control; values below 100% indicate inhibition. The associated SD was estimated using the delta method:

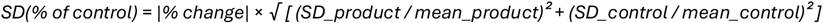

Where product counts fell below the detection limit (< 10 CFU/mL, based on a plating volume of 100 µL without dilution), the log_10_ change was reported as a minimum value (“> log_10_ change”) using the detection limit as a conservative substitute for the product mean. In figures, these values are indicated by a downward arrow (▼) with the label ‘BDL - below detection limit’. Results are presented per organism across the six tested concentrations (0.031%, 0.063%, 0.125%, 0.25%, 0.5%, and 1%).

## 3. Results: Effect of ThymoQuin^®^ on key species of the human intestine

To assess the susceptibility of selected human gut microbes to ThymoQuin^®^, four key bacterial species were tested *in vitro*: *A. muciniphila* (Fig.1), *K. pneumoniae* (Fig. 2), *L. plantarum* (Fig. 3), and *S. thermophilus* (Fig. 4). Minimum bactericidal concentration assays (MBCs) were performed using six different ThymoQuin^®^ concentrations, followed by incubation for 24 h under strictly anaerobic conditions, to get as close as possible to the conditions in the intestine. After incubation, bacterial suspensions were plated to determine the effects of ThymoQuin^®^ exposure on bacterial vitality relative to the untreated control.

**Figure 1:**
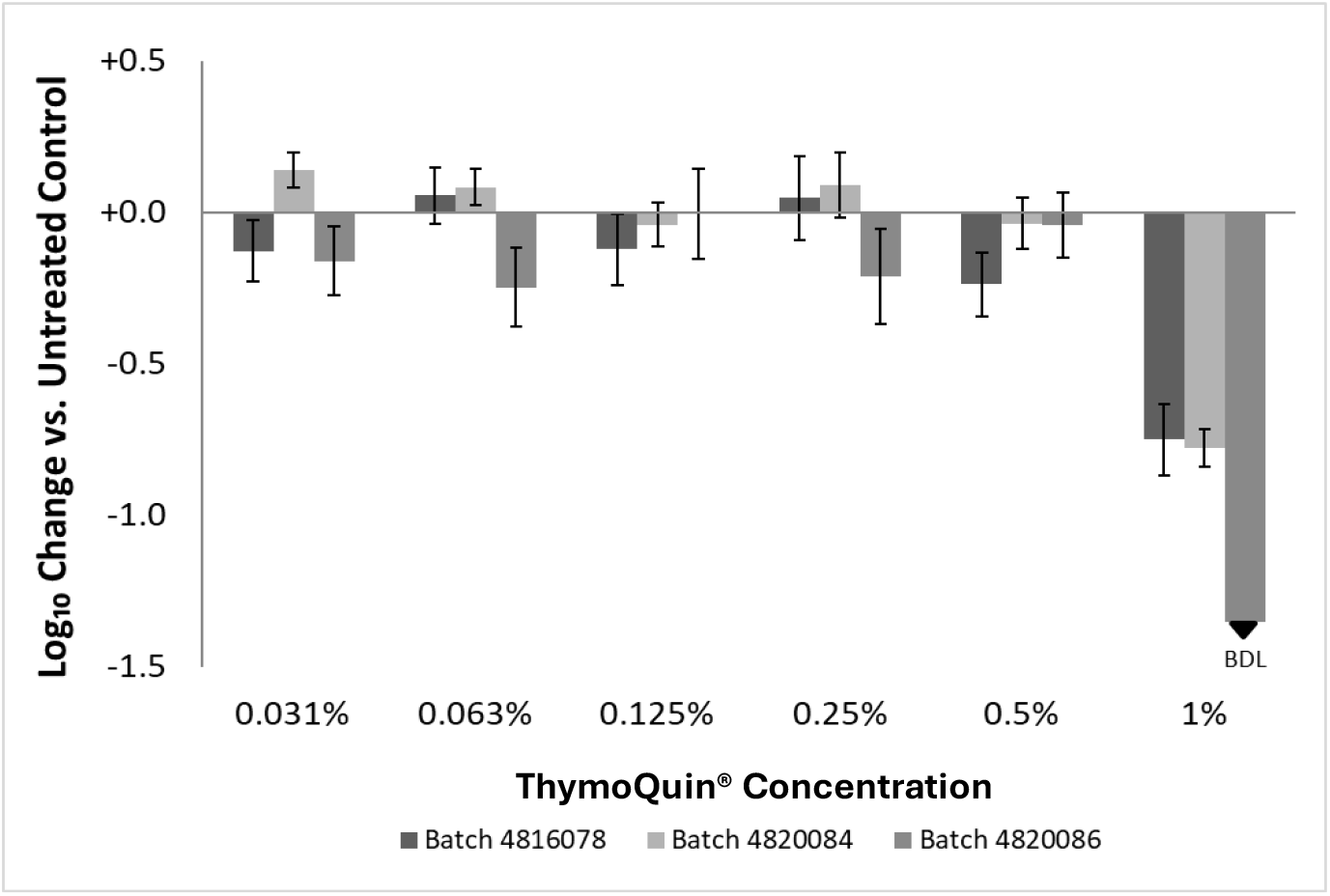
Effect of ThymoQuin^®^ on the vitality of *A. muciniphila*. *A. muciniphila* was exposed to six concentrations of ThymoQuin^®^ ranging from 0.031% to 1% for 24 h under anaerobic conditions. Three independent product batches, 4816078, 4820084, and 4820086, were tested (= 3 biolog. replicates, each in three technical replicates). Bars represent the mean log_10_ change in CFU/mL relative to the untreated control (log_10_ change = log_10_(CFU/mL treated) − log_10_(CFU/mL untreated control)), calculated from three technical replicates (n = 3). Error bars indicate the propagated standard deviation. Negative values indicate a reduction in viable counts relative to the untreated control; positive values indicate growth. The downward arrow (▼) indicates that viable counts in all replicates fell below the detection limit of 10 CFU/mL (corresponding to a minimum log_10_ reduction of 7.3), and the true value may therefore be lower than shown. The horizontal line at log_10_ change = 0 represents no change relative to the untreated control. The untreated control was prepared and incubated under identical conditions in the absence of the test substance.

**Figure 2:**
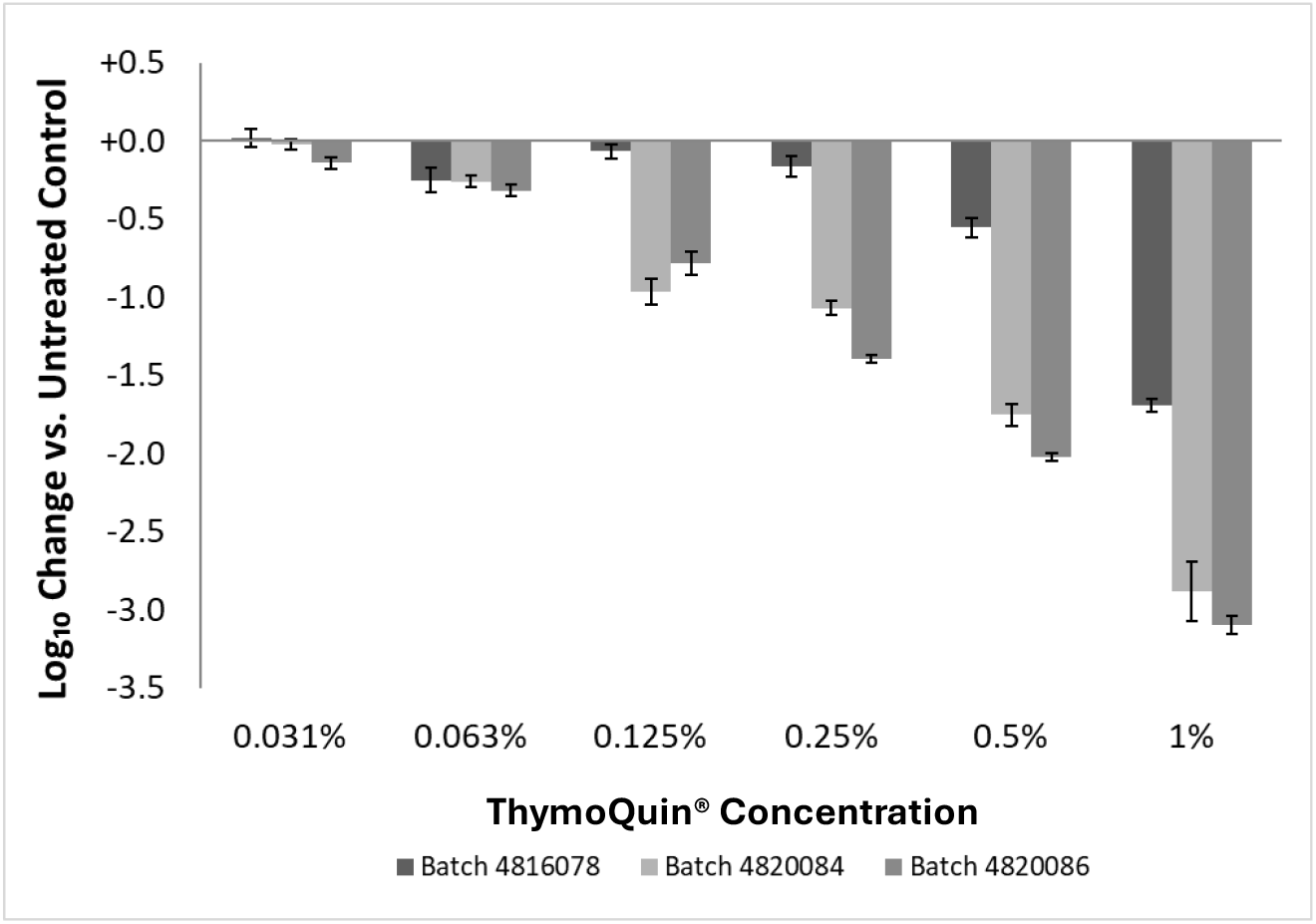
Effect of ThymoQuin^®^ on the vitality of *K. pneumoniae*. *K. pneumoniae* was exposed to six concentrations of ThymoQuin^®^ ranging from 0.031% to 1% for 24 h under anaerobic conditions. Three independent product batches, 4816078, 4820084, and 4820086, were tested (= 3 biolog. replicates, each in three technical replicates). Bars represent the mean log_10_ change in CFU/mL relative to the untreated control (log_10_ change = log_10_(CFU/mL treated) − log_10_(CFU/mL untreated control)), calculated from three technical replicates (n = 3). Error bars indicate the propagated standard deviation. Negative values indicate a reduction in viable counts relative to the untreated control; positive values indicate growth. The horizontal line at log_10_ change = 0 represents no change relative to the untreated control. The untreated control was prepared and incubated under identical conditions in the absence of the test substance.

**Figure 3:**
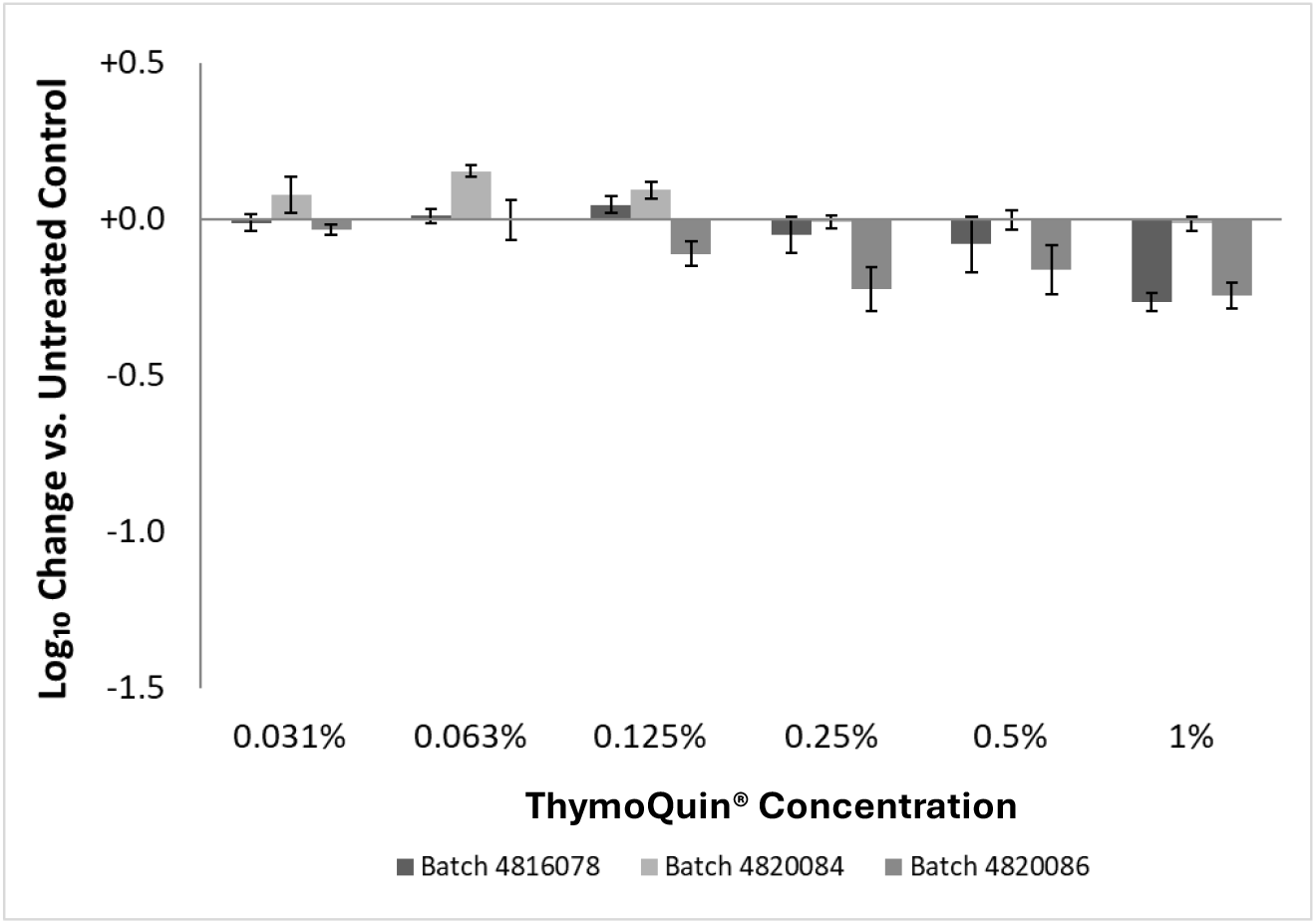
Effect of ThymoQuin^®^ on the vitality of *L. plantarum*. *L. plantarum* was exposed to six concentrations of ThymoQuin^®^ ranging from 0.031% to 1% for 24 h under anaerobic conditions. Three independent product batches, 4816078, 4820084, and 4820086, were tested (= 3 biolog. replicates, each in three technical replicates). Bars represent the mean log_10_ change in CFU/mL relative to the untreated control (log_10_ change = log_10_(CFU/mL treated) − log_10_(CFU/mL untreated control)), calculated from three technical replicates (n = 3). Error bars indicate the propagated standard deviation. Negative values indicate a reduction in viable counts relative to the untreated control; positive values indicate growth. The horizontal line at log_10_ change = 0 represents no change relative to the untreated control. The untreated control was prepared and incubated under identical conditions in the absence of the test substance.

**Figure 4:**
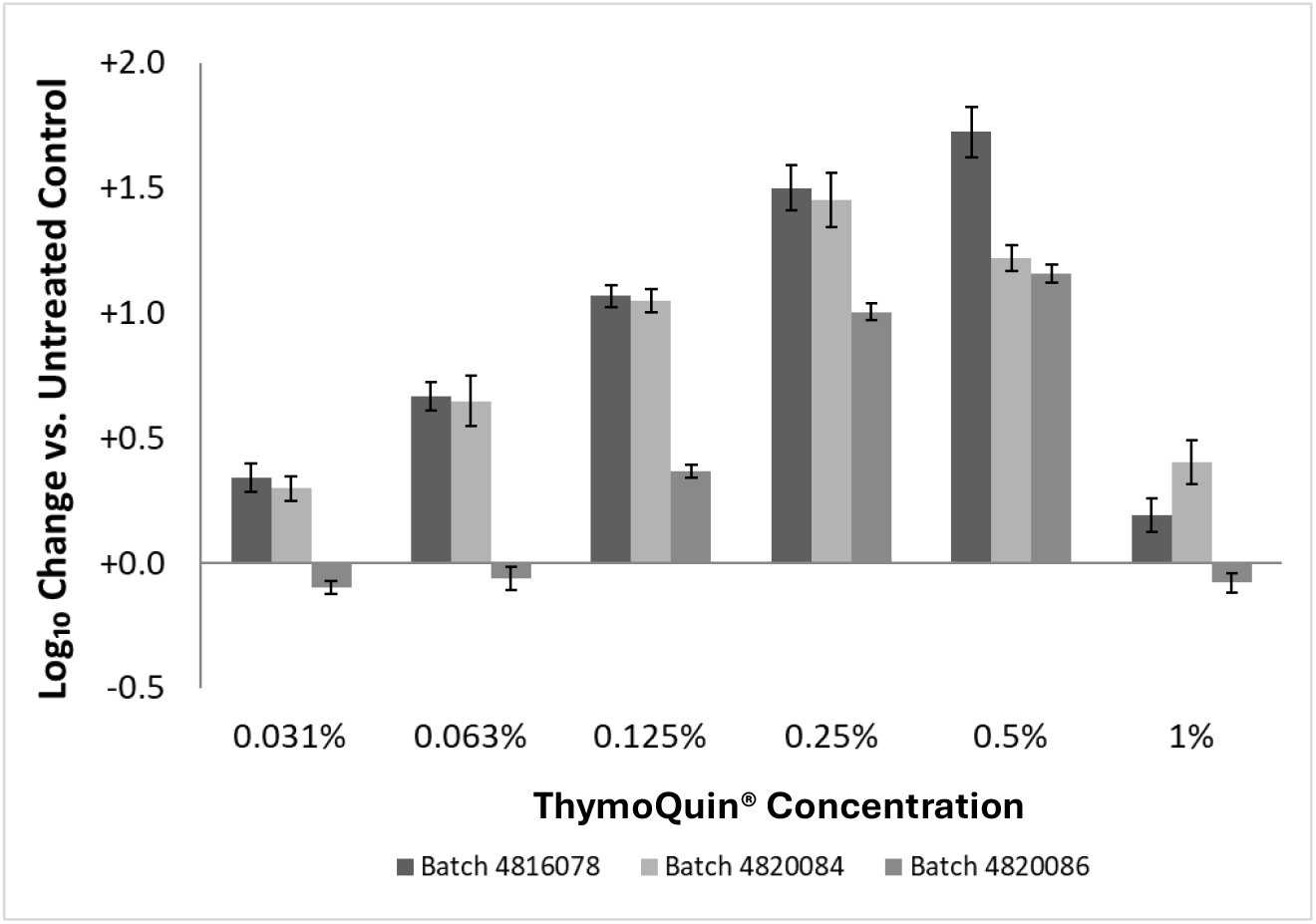
Effect of ThymoQuin^®^ on the vitality of *S. thermophilus*. *S. thermophilus* was exposed to six concentrations of ThymoQuin^®^ ranging from 0.031% to 1% for 24 h under anaerobic conditions. Three independent product batches, 4816078, 4820084, and 4820086, were tested (= 3 biolog. replicates, each in three technical replicates). Bars represent the mean log_10_ change in CFU/mL relative to the untreated control (log_10_ change = log_10_(CFU/mL treated) − log_10_(CFU/mL untreated control)), calculated from three technical replicates (n = 3). Error bars indicate the propagated standard deviation. Negative values indicate a reduction in viable counts relative to the untreated control; positive values indicate growth. The horizontal line at log_10_ change = 0 represents no change relative to the untreated control. The untreated control was prepared and incubated under identical conditions in the absence of the test substance.

### 3.1. Akkermansia muciniphila

The three ThymoQuin^®^ batches showed batch-dependent effects on the vitality of *A. muciniphila* (Fig. 1). At lower concentrations, particularly 0.031% and 0.063%, batches 4816078 and 4820084 increased bacterial vitality above the control level, suggesting a low stimulatory effect at low concentrations. In contrast, at the highest concentration tested, 1%, all batches markedly reduced vitality, indicating an inhibitory effect at elevated ThymoQuin^®^ concentrations. The response of *A. muciniphila* to ThymoQuin^®^ appeared to be primarily concentration-dependent, while batch-specific differences were comparatively minor. The data suggest that *A. muciniphila* largely tolerated concentrations up to 0.5%, whereas 1% ThymoQuin^®^ caused a marked reduction in bacterial vitality.

### 3.2 Klebsiella pneumoniae

*K. pneumoniae* (Fig. 2) depicted a concentration-dependent decrease in vitality following exposure to ThymoQuin^®^. At the lowest concentration, 0.031%, vitality remained close to the control level for batches 4816078 and 4820084, whereas batch 4820086 already showed a moderate reduction. With increasing concentration, bacterial vitality declined progressively, although the magnitude of the effect varied between batches. This batch-dependent variation was particularly evident at 0.125% and 0.25%, where batch 4816078 retained comparatively higher vitality, while batches 4820084 and 4820086 caused a pronounced reduction. At 0.5% and 1%, vitality was strongly reduced across all batches, approaching complete loss of viability at the highest concentration. Overall, *K. pneumoniae* was clearly sensitive to ThymoQuin^®^, with a pronounced concentration-dependent decline in vitality and evident batch-specific differences at intermediate concentrations.

### 3.3 *Lactiplantibacillus plantarum* subs. *Plantarum*

*L. plantarum* (Fig. 3) presented relatively stable vitality across the tested ThymoQuin^®^ concentration range, with most values remaining close to or above the control level. At lower concentrations, particularly 0.031% to 0.125%, batches 4820084 and 4820086 showed slightly increased vitality. At intermediate concentrations, 0.25% and 0.5%, vitality remained largely comparable to the control, with only minor batch-dependent variation. At 1%, a moderate reduction was observed for batches 4816078 and 4820086, whereas batch 4820084 remained close to the control level. Taken together, *L. plantarum* appeared to tolerate ThymoQuin^®^ well across the tested concentration range, showing only a moderate batch-dependent reduction at the highest concentration.

### 3.4 Streptococcus thermophilus

MBCs with *S. thermophilus* (Fig. 4) resulted in a pronounced concentration-dependent increase in vitality following exposure to ThymoQuin^®^. At low concentrations, 0.031% and 0.063%, vitality showed already dramatically increased values of up to 460% compared to the control. An even stronger stimulatory effect was observed at 0.125% with up to 1170% vitality, and this effect became particularly pronounced at 0.25% and 0.5%, where vitality values increased about 50-fold above the control level. The magnitude of this response differed between batches, with batch 4816078 showing the strongest increase, especially at 0.25% and 0.5%. At 1%, vitality declined markedly compared with the mid-range concentrations, although values did not indicate the same strong stimulatory response observed at 0.25–0.5%.

### 3.5 Analysis of the results by ThymoQuin^®^ batches

Across all three batches (Fig. 5), ThymoQuin^®^ exerted markedly different effects depending on the bacterial strain and concentration tested. *S. thermophilus* consistently showed the highest viability ratios, frequently exceeding 10^3^ at mid-range concentrations (0.125%–0.5%), suggesting a pronounced growth-promoting or stress-tolerant response. *A. muciniphila* and *L. plantarum* generally maintained viability ratios near or above 10^2^ across most concentrations, indicating limited inhibition. In contrast, *K. pneumoniae* demonstrated a concentration-dependent reduction in viability, with ratios dropping below 10^1^ at 0.5%–1% in batches 4816084 (Fig. 5A) and 4816086 (Fig. 5C) and reaching values below 10^0^ (i.e., below the untreated control level) at 1% in batch 4816084 (Fig. 5A) and batch 4816086 (Fig. 5C). Batch 4816078 (Fig. 5B) showed a broadly similar pattern, although the inhibitory effect on *K. pneumoniae* appeared slightly less pronounced at the highest concentrations. Overall, results were qualitatively consistent across batches, supporting the reproducibility of the observed effects.

**Figure 5.**
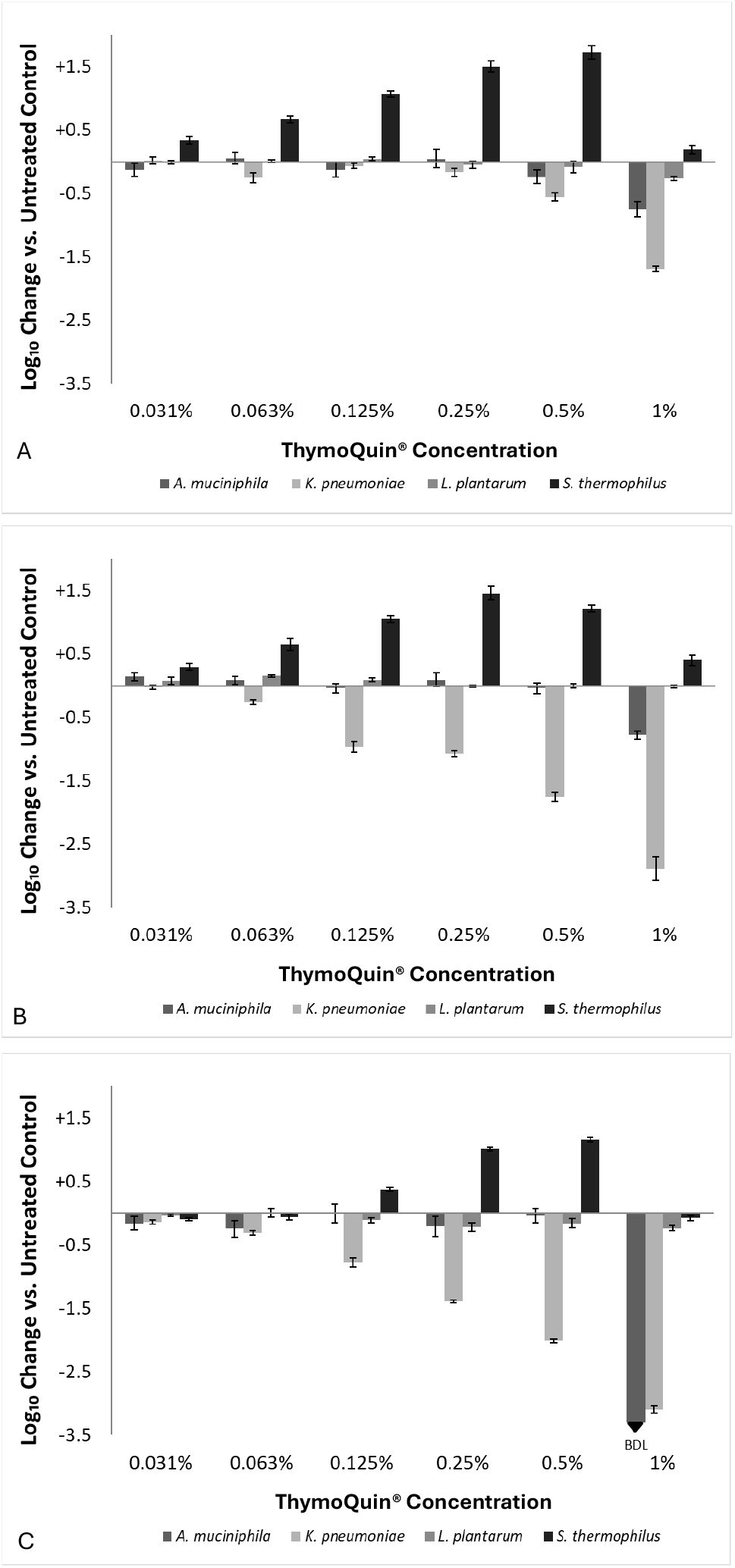
Effect of ThymoQuin^®^ on the growth of four microorganisms at increasing concentrations, shown per batch. Panels A–C show results for Batch 4816078, 4820084, and 4820086, respectively. Each panel displays the log_10_ change in CFU/mL relative to the untreated control (log_10_ change = log_10_(CFU/mL treated) − log_10_(CFU/mL untreated control)) for *A. muciniphila, K. pneumoniae, L. plantarum*, and *S. thermophilus*. Bars represent the mean of three technical replicates (n = 3). Error bars indicate the propagated standard deviation. Negative values indicate a reduction in viable counts relative to the untreated control; positive values indicate growth. The downward arrow (▾) indicates that viable counts in all replicates fell below the detection limit of 10 CFU/mL, and the true value may therefore be lower than shown. The horizontal line at log_10_ change = 0 represents no change relative to the untreated control. The untreated control was prepared and incubated under identical conditions in the absence of the test substance.

## 4. Discussion

Black seed oil, obtained from *Nigella sativa* seeds, has gained scientific interest because it contains both nutritional lipids and bioactive compounds. Reported properties include antioxidant, anti-inflammatory, immunomodulatory, antimicrobial, metabolic, and gastrointestinal effects, which are often linked to thymoquinone, one of the major bioactive constituents of the volatile fraction. However, black seed oil is not a single-compound substance; it contains fatty acids, volatile compounds, phenolics, sterols, and quinone derivatives, which may all contribute to its biological activity [23].

A key issue when comparing studies is the large variability between *N. sativa* preparations. Conventional black seed oil, thymoquinone-rich or standardized oils, isolated thymoquinone, seed powder, extracts, and essential oil can differ substantially in chemical composition. Fixed oils are mainly composed of fatty acids such as linoleic, oleic, and palmitic acids, whereas the volatile fraction contains compounds such as thymoquinone, p-cymene, thymohydroquinone, and thymol. Therefore, biological effects may depend strongly on the preparation used, the thymoquinone content, extraction method, origin, processing, and storage conditions [24].

In the frame of this study, ThymoQuin^®^, black cumin seed oil, developed by Trinutra, was tested towards its effects on different key bacterial species present in the human gut. The present findings indicate that ThymoQuin^®^ exerts a concentration-dependent and species-specific effect on the tested bacterial strains. Across all evaluated batches, ThymoQuin^®^ promoted the growth of the probiotic strain *S. thermophilus* over a broad concentration range, while *L. plantarum* was largely preserved at low to intermediate concentrations. In contrast, the opportunistic pathogen *K. pneumoniae* showed a progressive reduction in viability with increasing ThymoQuin^®^ concentrations. This pattern suggests a selective antimicrobial profile, in which beneficial or probiotic-associated bacteria may be maintained or stimulated within a defined concentration window, whereas a pathogenic strain is inhibited.

The strongest probiotic-promoting effect on the members of the gut microbiome was observed, when *S. thermophilus* was exposed to different ThymoQuin^®^ concentrations. ThymoQuin^®^ concentrations between 0.031% and 0.5% increased *S. thermophilus* growth, with the most pronounced stimulation observed at 0.25% and 0.5%. In contrast, the highest tested concentration, 1%, resulted in a marked collapse of *S. thermophilus* growth, indicating that the beneficial effect is limited to a defined concentration range and that excessive concentrations may become inhibitory even for probiotic species. *L. plantarum* showed a more stable but less strongly stimulated response, with growth largely preserved between 0.031% and 0.5%, while visible stress or reduced recovery was observed at 1%. A similar dose-dependent safety aspect was observed for *A. muciniphila*. Across the different batches, ThymoQuin^®^ had little or no consistent growth-promoting or inhibitory effect on *A. muciniphila* at low to intermediate concentrations, whereas the highest concentration showed a pronounced bactericidal effect. This finding is relevant because *A. muciniphila* is a mucus-associated commensal bacterium with context-dependent effects on host health. It has been linked to beneficial metabolic and barrier-related outcomes, but excessive or inappropriate modulation of mucin-degrading bacteria may also have unfavorable consequences under certain conditions [18]. Therefore, the lack of major effects on *A. muciniphila* within the lower concentration range may be interpreted as favorable, whereas the bactericidal effect at 1% further supports the need to avoid overdosing [25]. These results suggest that concentrations below 1% may be compatible with selected probiotic bacteria under the present experimental conditions, whereas 1% appears too high for a selectively probiotic-supportive application.

Importantly, the response of *K. pneumoniae* differed markedly from that of the probiotic strains. Even at the lowest concentration tested, ThymoQuin^®^ produced a mild reduction in *K. pneumoniae*, and this inhibitory effect increased progressively with dose. At 0.25% and 0.5%, *K. pneumoniae* was strongly reduced, while 1% resulted in near-total inhibition. Taken together, these data suggest that the most relevant concentration window lies between 0.125% and 0.5%, where *S. thermophilus* was strongly stimulated, *L. plantarum* was largely preserved, and *K. pneumoniae* was substantially inhibited. Among these concentrations, 0.25% may represent a particularly balanced dose, because it combined strong stimulation of *S. thermophilus*, preservation of *L. plantarum*, and marked inhibition of *K. pneumoniae*, without reaching the toxicity-associated pattern observed at 1%.

These *in vitro* findings are consistent with a previously published randomized, double-blind, placebo-controlled clinical study investigating ThymoQuin^®^ black cumin seed oil supplementation in endurance runners [26]. In that study, 37 runners consumed 500 mg/day of ThymoQuin^®^ 3% or placebo for four weeks, spanning three weeks before and one week after a half-marathon or marathon event. Compared with placebo, ThymoQuin^®^ supplementation was associated with significantly fewer upper-respiratory tract complaints, improved global mood state, lower salivary cortisol, higher microbiome composite score, and a 66% higher relative abundance of *S. thermophilus* after supplementation.

The clinical observation of increased *S. thermophilus* abundance after ThymoQuin^®^ supplementation is particularly relevant to the present study, because *S. thermophilus* was also the bacterial species that showed the strongest growth-promoting response *in vitro*. While the clinical trial measured changes in fecal microbiome composition in humans and the present study assessed direct bacterial responses under controlled *in vitro* conditions, both datasets point in the same direction: ThymoQuin^®^ may create conditions that favor *S. thermophilus* abundance or growth. This convergence strengthens the biological plausibility of a selective microbiome-modulating effect, although the mechanisms cannot be established from the current data alone.

The clinical study [26] also reported broader host-related outcomes, including 62% fewer upper-respiratory tract complaints, an 11% improvement in global mood state, an 8-point higher microbiome composite score, and 44% lower salivary cortisol compared with placebo after four weeks. These findings suggest that ThymoQuin^®^ may influence not only specific microbial taxa but also host parameters related to stress physiology, immune vigilance, and subjective well-being. However, these outcomes should be interpreted cautiously, because the trial was conducted in a specific population of endurance athletes exposed to exercise-induced physiological stress. Nevertheless, the concordance between the present *in vitro* findings and the clinical data by Talbott and Talbott [26] supports the translational relevance of the experimental model.

This is important because in vitro microbiological models are often criticized for not fully reproducing the complexity of the human gastrointestinal environment. Indeed, host metabolism, immune signaling, bile acids, intestinal transit, and microbial cross-feeding cannot be completely replicated in simplified laboratory systems. However, *in vitro* gut fermentation and microbiota models are widely used to study microbial modulation under controlled and reproducible conditions and are considered valuable complementary tools to human studies [27, 28]. Thus, the present data does not prove full equivalence between *in vitro* and *in vivo* conditions, but they provide supportive evidence that carefully designed *in vitro* assays can identify microbiological effects with *in vivo* relevance. In the case of ThymoQuin^®^, the matching *S. thermophilus* response strengthens the interpretation that the observed growth-promoting effect is not merely an artificial laboratory finding but may reflect a relevant microbiome-modulating property of thymoquinone-rich black seed oil [29, 30].

Overall, the present results support the concept that ThymoQuin^®^ has a hormetic dose-dependent and selective antimicrobial profile, which is generally characterized by stimulation at low to moderate doses and inhibition at higher doses [31]. This biphasic response supports the interpretation that ThymoQuin^®^ may act as a microbial modulator within an optimal concentration window, while excessive exposure becomes inhibitory. Hormetic dose responses have been described across biological systems and are considered a relevant concept for interpreting non-linear effects of bioactive compounds [32]. Concentrations between 0.125% and 0.5% appear most promising, as they combined probiotic compatibility, pronounced *S. thermophilus* stimulation, and substantial inhibition of *K. pneumoniae*. The parallel with the published *in vivo* study, in which ThymoQuin^®^ supplementation increased *S. thermophilus* abundance and improved several immune- and stress-related outcomes, provides supportive external evidence for the relevance of the observed *S. thermophilus* response [26]. Future studies should include statistical analysis of batch variability, additional probiotic and pathogenic species, mixed-community microbiome models, and mechanistic assays to clarify whether ThymoQuin^®^ acts through direct bacterial growth modulation, selective antimicrobial pressure, altered metabolic availability, or host-mediated pathways.

